# Dengue virus protease activity modulated by dynamics of protease cofactor

**DOI:** 10.1101/2020.11.11.378646

**Authors:** W H Lee, W Liu, J Fan, D Yang

## Abstract

The viral protease domain (NS3pro) of dengue virus is essential for virus replication and its cofactor NS2B is indispensable for the proteolytic function. Although several NS3pro-NS2B complex structures have been obtained, the dynamic property of the complex remains poorly understood. Using NMR relaxation techniques, here we found that NS3pro-NS2B exists in both closed and open conformations which are in dynamic equilibrium on a sub-millisecond timescale in aqueous solution. Our structural information indicates that the C-terminal region of NS2B is disordered in the open conformation but folded in the closed conformation. Using mutagenesis, we showed that the closed-open conformational equilibrium can be shifted by changing NS2B stability. Moreover, we revealed that the proteolytic activity of NS3pro-NS2B correlates well with the population of the closed conformation. Our results suggest that the closed-open conformational equilibrium can be used by both nature and man to control the replication of dengue virus.

**Statement of Significance:** The dengue virus protease is an attractive target for drug development against dengue fever as it is essential for virus replication. However, its structure-based drug development has been unsuccessful due to the shallow substrate-binding pocket. The study presented here demonstrates for the first time that the protease activity can be reduced dramatically by shifting the closed-open conformational equilibrium of the protease in complex with its cofactor from the majority of a closed conformation to the majority of an open conformation. Moreover, our work clarifies the structure of the open conformation which has been elusive for a long time. Our results also suggest an alternative method for designing protease inhibitors based on the closed-open conformational equilibrium.

## Introduction

Dengue fever is a disease that is caused by the dengue virus, which is transmitted by the mosquito species *Aedes aegypti*. It has been estimated that up to 4 billion people are at risk of dengue fever globally(1), with as many as 400 million new infections annually(2). Symptoms of dengue fever include fever, headache, muscle and joint pains, nausea, vomiting, swollen glands and rashes(3). More severe cases manifest as dengue haemorrhagic fever or dengue shock syndrome and can be fatal(4). The dengue vaccine Dengvaxia has been approved recently, and a few more are undergoing clinical trials. However, Dengvaxia can cause antibody-dependent enhancement (ADE) and is therefore not recommended for people who has never been infected with dengue before(5). Meanwhile, there are no approved drugs for treating dengue yet. It is therefore of great importance and urgency to understand more about the dengue virus, so that effective treatments can be developed against it.

The genome of the dengue virus consists of a positive-sense single-stranded RNA of about 11 kb long(6). It contains an open reading frame that encodes for a single polyprotein containing three structural proteins and seven non-structural proteins(3, 7). For each of the proteins to perform their respective functions, the polyprotein needs to be cleaved at various sites by both the host proteases and the viral protease(8). If the proteolytic processing of the viral polyprotein can be disrupted, the dengue virus will no longer be able to replicate and propagate, effectively stopping the progression of dengue fever on its tracks. This makes the viral protease an attractive target for drug development. In order to do so, sufficient structural and dynamic information of the dengue protease is required.

The N-terminal protease domain of non-structural protein 3 (NS3pro) is a chymotrypsin-like serine protease responsible for the cleavage of the polyprotein at various sites(9-12). NS3pro also cleaves within some viral proteins like the capsid (C) protein(13), NS2A(14), NS4A(15) and even NS3 itself (16, 17). NS3pro is unable to fold correctly by itself and requires the hydrophilic region of non-structural protein 2B (NS2B) as a cofactor for its correct folding and protease activity(9, 18, 19). The N-terminal 18 amino acids of the hydrophilic region of NS2B (residues 49 to 66) allow NS3pro to adopt its correct structure by contributing a β-strand to complete the β-barrel of NS3pro(18). However, this region of NS2B is still insufficient for the protease activity of NS3pro(18, 19). The remaining C-terminal portion of the hydrophilic region of NS2B (residues 67 to 95) contributes a β-hairpin that completes the substrate-binding pocket of NS3pro and is therefore necessary for its activity(20).

Existing crystal and solution structures of the NS2B-NS3pro complex show that NS2B can adopt drastically different conformations(18-26). In the absence of an inhibitor, the C-terminal region of NS2B turns away from the catalytic triad of NS3pro in the crystal state and is denoted as the open conformation (PDB ID: 2FOM)(18). In the solution state, no inhibitor-free structure is available, probably due to the dynamic nature of NS2B(27, 28). On the other hand, in the presence of an inhibitor, the C-terminal region of NS2B is in close proximity to the catalytic triad of NS3pro and completes the substrate-binding pocket (PDB ID: 2FP7)(18). This is denoted as the closed conformation, which is the enzymatically active form. Intriguingly, previous NMR studies have revealed that the closed conformation is the dominant state even in the absence of an inhibitor though the exact structure is unknown(28). In the crystal structure of NS2B-NS3pro with aprotinin (PDB ID: 3U1J), the electron density for the whole C-terminal region of NS2B (residues 70 to 95) was missing(20), suggesting a highly dynamic nature of this region. The presence of both the open and closed conformations suggests the protein can undergo conformational exchange, further supporting the dynamic nature of the protein. So far, no detailed studies on the dynamics NS2B-NS3pro complex have been reported. Moreover, the relationship between the dynamics and protease activity of NS2B-NS3pro is still totally unclear.

In this study, we investigated the role of NS2B dynamics in the catalytic activity of NS2B-NS3pro using NMR relaxation techniques, mutagenesis, and activity assay. Structural information of the elusive open conformation was revealed by NMR dynamics experiments, while the activity level of the dengue protease was found to be correlated to the population size of the closed conformation.

## Materials and Methods

### Cloning of DENV2 NS2B and NS3pro

The codon-optimised gene encoding DENV2 NS3pro (residues 14–185, based on the accession number ARO84675.1) was synthesized by GenScript and ligated into pSKDuet01 using the sites NcoI and XhoI. DENV2 NS2B (residues 48–100) was assembled from four primers using PCR, after which an N-terminal 6xHis-Smt3 tag was added by PCR. The resulting PCR product was ligated into pSKBAD2 using the sites NdeI and HindIII. pSKDuet01 and pSKBAD2 were gifts from Hideo Iwai (Addgene plasmids # 12172 and # 15335)(29).

Point mutants of DENV2 NS2B-NS3pro were generated through site-directed mutagenesis. The mutant inserts were generated in a first PCR step as two fragments. The two fragments were then linked by a second PCR step to generate a single fragment. PCR products of NS2B mutants were ligated into pSKBAD2 using the sites NdeI and HindIII, while PCR products of NS3pro mutants were ligated into pSKDuet01 using the sites NcoI and XhoI.

### Expression of DENV2 NS2B-NS3pro complex

BL21 (DE3) cells with both plasmids containing 6xHis-Smt3-NS2B and NS3pro were grown in Lysogeny Broth (LB) medium with 100 µg/ml ampicillin and 25 µg/ml kanamycin at 37 °C and 200 rpm until an OD_600_ of 0.6 was reached. Expression of 6xHis-Smt3-NS2B was induced with 2 g/L arabinose for one hour at 37 °C and 200 rpm, after which 0.1 mM isopropyl β-D-1-thiogalactopyranoside (IPTG) was added to induce the expression of NS3pro at 30 °C and 200 rpm for three hours.

For the expression of isotopically labelled NS2B complexed with unlabelled NS3pro, a protocol derived from Muona et al. was used(30). BL21 (DE3) cells with both plasmids containing 6xHis-Smt3-NS2B and NS3pro were grown in the same LB medium and condition as that for the non-labelled protein until an OD_600_ of 0.6 was reached. The cells were harvested by centrifugation and then transferred to M9 medium. After growing for one hour, 2 g/L arabinose was added to induce the expression of 6xHis-Smt3-NS2B for three hours at 37 °C and 200 rpm. Subsequently, the cells were collected by centrifugation. After washing the cells with Terrific Broth (TB), the cell pellet was resuspended in TB medium containing 100 µg/ml ampicillin, 25 µg/ml kanamycin. Finally, 0.1 mM IPTG was added to induce the expression of NS3pro for three hours at 30 °C and 200 rpm.

### Purification of DENV2 NS2B-NS3pro complex

The cells were harvested by centrifugation and lysed by sonication. The protein was purified from the supernatant of the lysate using a 5 ml HisTrap HP column (GE Healthcare). The Smt3 tag was cleaved off before further purification by a HiLoad 16/600 Superdex 75 size exclusion column (GE Healthcare) in NMR buffer (20 mM sodium phosphate pH 6.5, 10 mM sodium chloride, 1 mM ethylenediaminetetraacetic acid (EDTA)). Aliquots were prepared and stored at −80 °C for activity assays.

### Activity Assay of DENV2 NS2B-NS3pro

Activity assays were performed in 100 µl reaction volumes on a 96-well plate as described in a previous study(31). Briefly, 10 nM NS2B-NS3pro was incubated with the substrate benzoyl-Nle-Lys-Arg-Arg-7-amino-4-methylcoumarin (Bz-nKRR-AMC) (Genscript) at concentrations varying from 2 to 200 µM. The assay buffer used was 50 mM Tris (pH 8.5), 20% glycerol (v/v) and 1 mM 3-[(3-cholamidopropyl)dimethylammonio]-1-propanesulfonate (CHAPS). The fluorescence was then monitored in a Tecan Infinite 200 PRO microplate reader at 25 °C, with excitation and emission wavelengths of 380 nm and 450 nm respectively. Fluorescence readings were converted to AMC concentrations using a standard AMC curve. Data points from the activity assay were fitted to the Michaelis-Menten equation(32).

### Circular dichroism (CD) spectroscopy

CD spectra were recorded on samples containing 10 µM protein in the NMR buffer using a Jasco J-1100 CD spectrophotometer at 20 °C.

### NMR spectroscopy

NMR samples were prepared at concentrations of 0.5–0.7 mM protein in the NMR Buffer plus 5% D_2_O. All NMR experiments were performed on a Bruker AVANCE 800 MHz spectrometer equipped with a cryoprobe at 25 °C. NMR data were processed using NMRPipe and analysed using SPARKY(33).

3D HNCOCA, HNCA and ^15^N-edited NOESY-HSQC spectra were acquired on ∼0.7 mM of ^13^C,^15^N-labelled NS2B in complex with unlabelled NS3pro. Heteronuclear ^15^N nuclear Overhauser effects (NOEs) were measured on a ^15^N-labelled NS2B complexed with unlabelled NS3pro (∼0.5 mM) from two sets of data obtained in the presence and absence of proton saturation. In the measurements, the recycle delay was 8 s and the proton saturation time was 4 s. The ^15^N NOE values were derived from ratios of the peak intensities with and without proton saturation(34).

CPMG relaxation dispersion experiments were performed on a ^15^N-labelled NS2B-unlabelled NS3pro complex (∼0.5 mM) using a pulse sequence established previously(35). A constant CPMG delay (*T*_*CPMG*_) of 20 ms was used for each CPMG experiment. The CPMG frequencies (*v*_*CPMG*_) used were 50, 100, 150, 200, 250, 300, 400, 500, 600, 700, 800, 900, 1000, 1200, 1400, 1600 and 2000 Hz. The CPMG experiment at *v*_*CPMG*_ = 100 Hz was performed twice to estimate the experimental error. The total experimental time was about 18 hours.

^15^N CEST experiments were performed on a ^15^N-labelled NS2B-unlabelled NS3pro complex (∼0.5 mM) using two weak RF fields of 13 and 26 Hz, with a CEST mixing time of 0.5 s. At the higher RF field, 27 ^1^H-^15^N HSQC spectra were collected with ^15^N carrier frequencies from 107 to 133 ppm at 1 ppm intervals. On the other hand, at the lower RF frequency, 38 ^1^H-^15^N HSQC spectra were collected with ^15^N carrier frequencies from 106 to 135 ppm at 0.5 ppm intervals between 115 and 124 ppm, and 1 ppm intervals otherwise. The total experimental time was 44 hours. For peaks with ^15^N chemical shift greater than 120 ppm, data points before 111 ppm were used to estimate the error. On the other hand, for peaks with ^15^N chemical shifts smaller than 120 ppm, data points after 129 ppm were used to estimate the error.

### Analyses of NMR data

Backbone ^15^N and ^1^HN assignments of NS2B were achieved using 3D HNCA and HNCOCA spectra because of a small number of residues. In the cases where ambiguities occurred due to similar ^13^Cα chemical shifts, ^15^N-edited NOESY-HSQC was used.

To obtain conformational exchange parameters and chemical shifts of the minor state, relaxation dispersion and CEST data of residues displaying obvious relaxation dispersion were simultaneously fitted to a 2-state exchange model (N *⇌* I, where N and I represent the major native state and minor non-native state respectively) as described previously(36). Briefly, the data for each residue (j) were fitted to estimate individual exchange rate (*k*_*ex*_*(j)*), population (*p*_*I*_*(j)*), chemical shift (*δ*_*I*_*(j)*), intrinsic transverse relaxation rate (R_2_*(j)*), and longitudinal relaxation rate (R_1_*(j)*). From these estimations, initial values of the fitting parameters were determined roughly. Next, the data for all the residues were fitted globally to extract global kinetics parameters (exchange rate and population) and residue-specific parameters (chemical shifts and relaxation rates). In the fitting, the R_2_ and R_1_ values for each residue were assumed to be independent of conformational states. Error estimation also followed the previous method (36).

### Chemical shift perturbation

For a given residue, the chemical shift perturbation was calculated using the equation,

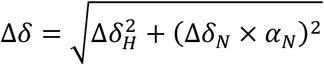

where *Δδ*_*H*_ (*Δδ*_*N*_) is the difference of ^1^H (^15^N) chemical shifts between the WT protein and its mutant, and *α*_*N*_ is the scaling factor for the ^15^N chemical shift difference(37). A value of 0.17 was used for *α*_*N*_ in this study.

## Results

### Dynamics of NS2B in NS2B-NS3pro complex on ns–ps timescale

NS2B-NS3pro complex consists of 225 amino acid residues. A previous study showed that about 50% of the NS2B ^1^H-^15^N correlations (peaks) are unavailable for providing dynamic information mainly due to peak overlap(38). To overcome this overlapping problem, we obtained a sample of ^13^C,^15^N-labelled NS2B (S48-R100) complexed with unlabelled NS3pro using a sequential expression approach described previously(30). Except for S48 and D81, backbone ^1^H-^15^N correlations for all other residues were observable in the HSQC spectrum with good peak dispersion (Fig. 1). The assignments were achieved using triple resonance experiments.

**Figure 1.**
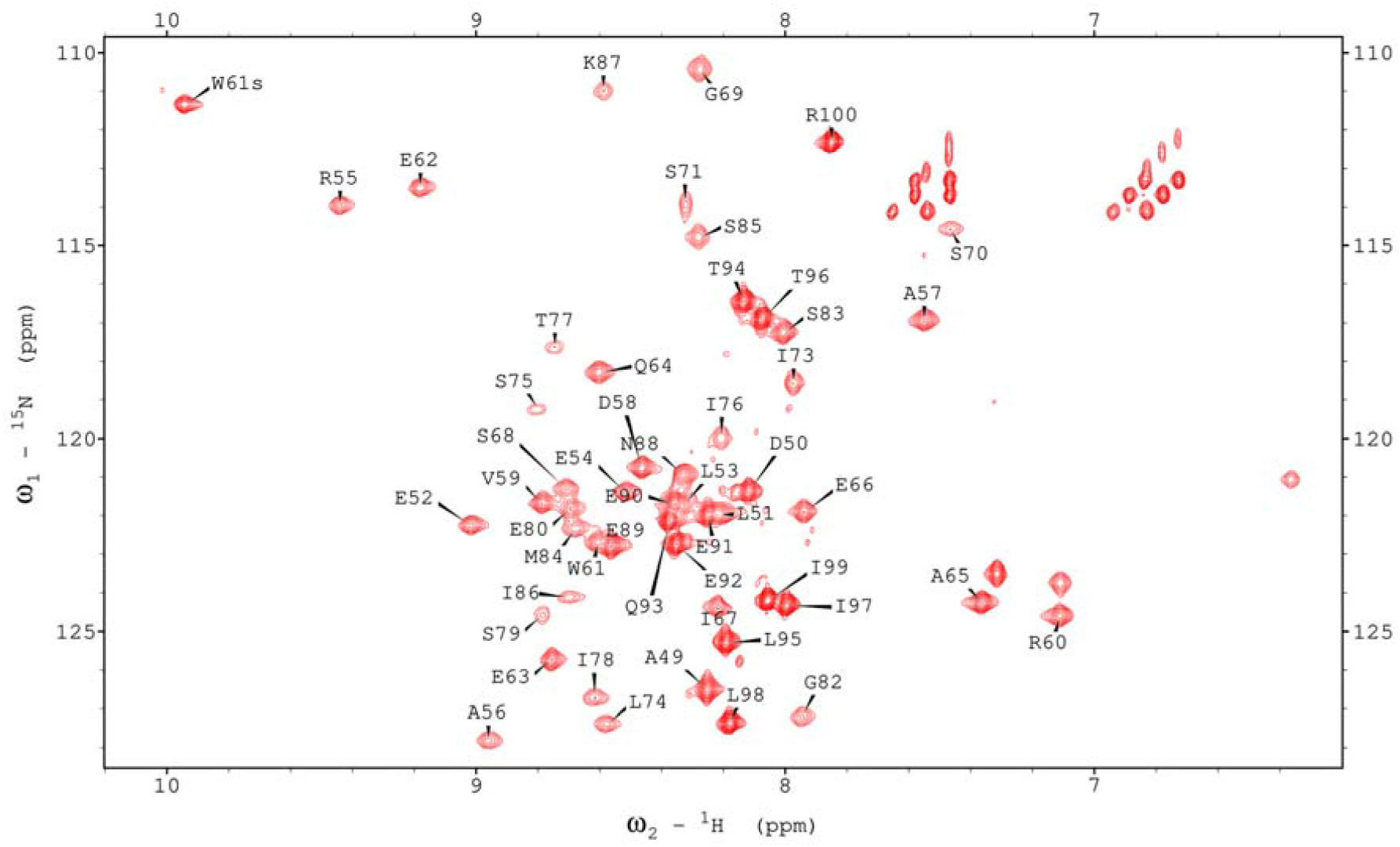
HSQC spectrum of ^15^N-labelled DENV2 NS2B complexed with unlabelled NS3pro, with peak assignments included. The peaks of W61s (sidechain HεNε), G82 and K87 are aliased by 20 ppm in the ^15^N dimension.

The rigidity of NS2B in the wild-type complex was probed through the measurement of heteronuclear ^15^N NOE (HetNOE) values. High HetNOE values (>0.5) were observed for residues E52–Q64, E66–S68 and I73–K87 of NS2B, and relatively low values (<0.45) for C-terminal region (N88–R100), N-terminal end (A49–L51), Q64, A65, E66, G69 and S70 (Fig. 2). The result indicates that the C-terminal region N88–R100 and N-terminal end are flexible on the nanosecond to picosecond timescale. This is consistent with previous X-ray crystallography studies, which showed that the C-terminal region is undetectable in both closed and open conformations due to high flexibility and lack of regular helical or β-strand structure (18, 20, 21). In the closed conformation of NS2B-NS3pro complex, NS2B consists of four short β-strands (β1: D50-A57, β2: A65-S68, β3: I73-I78, β4: S83-I86) and three loops (loop1: D58-Q64, loop2: G69-P72, loop3: S79-G82) between the strands. Interestingly, most of residues in these loops are as rigid as those residues in the β-strands. Previous NMR studies found that the residues in the β-strands of NS2B have similar rigidity as those in the β-strands of NS3pro(28). In the open conformations of NS2B-NS3pro complexes which were solved previously by crystallography, except for the first β-strand (D50–A57), all other regions have no strong interactions such as H-bonding with NS3pro and thus adopt different conformations and orientate differently in different structures. For such open conformations, the region of E63–K87 should be significantly more flexible than the first β-strand (D50–A57). In fact, except a few residues in loop1, β2 and loop2 (Q64, A65, E66, G69 and S70), all other residues within E52-K87 have similar rigidity. Therefore, most residues in the region of E52–K87 should be in contact with NS3pro, and the wild-type NS2B-NS3pro complex is mainly in the closed conformational state.

**Figure 2.**
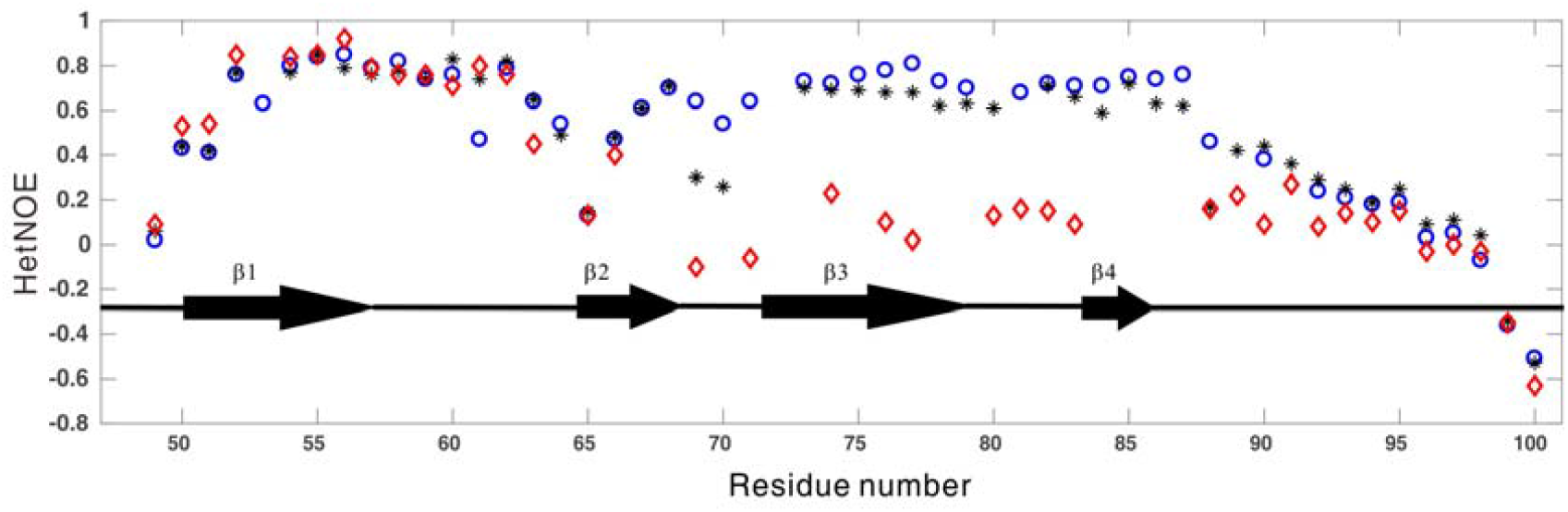
Heteronuclear NOE values of NS2B residues for the wild-type (black asterisks), the NS2B M84P mutant (red diamonds) and the NS2B I73C+NS3pro L115C disulfide bond mutant (blue circles). The secondary structure of NS2B is included within the plot.

### Dynamics of NS2B in NS2B-NS3pro complex on ms–μs timescale

To probe conformational exchange on ms–µs timescale, we carried out relaxation dispersion (RD) experiments. Residues located in the C-(A49–E62) and N-terminal (E89–R100) regions displayed nearly no ^15^N RD with R_ex_ values smaller than 1 s^-1^ (Fig. 3a, g). On the other hand, 21 out of 24 residues with available assignments in the region of E63–N88 exhibited significant RD with R_ex_ values larger than 3 s^-1^ (Fig. 3c, e). Here, R_ex_ is defined as R_2_eff(2000) - R_2_^eff^(50), where R_2_^eff^(2000) and R_2_^eff^(50) were the relaxation rates measured at CPMG fields of 2000 and 50 Hz, respectively. The RD data indicate that at least one “invisible” minor state (I) is in dynamic equilibrium with the observed major native state (N) on millisecond timescale for the region of E63–N88. In order to examine if slow conformational exchange processes on sub-second timescales exist, chemical exchange saturation (CEST) experiments were also performed. No residues displayed two obvious dips, indicating that the conformational exchange is too fast to be detected by CEST. To obtain structural information of the “invisible” state and kinetics parameters for the conformational exchange process, a 2-state model (N ⇌ I) was used to fit both the CEST and RD data simultaneously.

**Figure 3.**
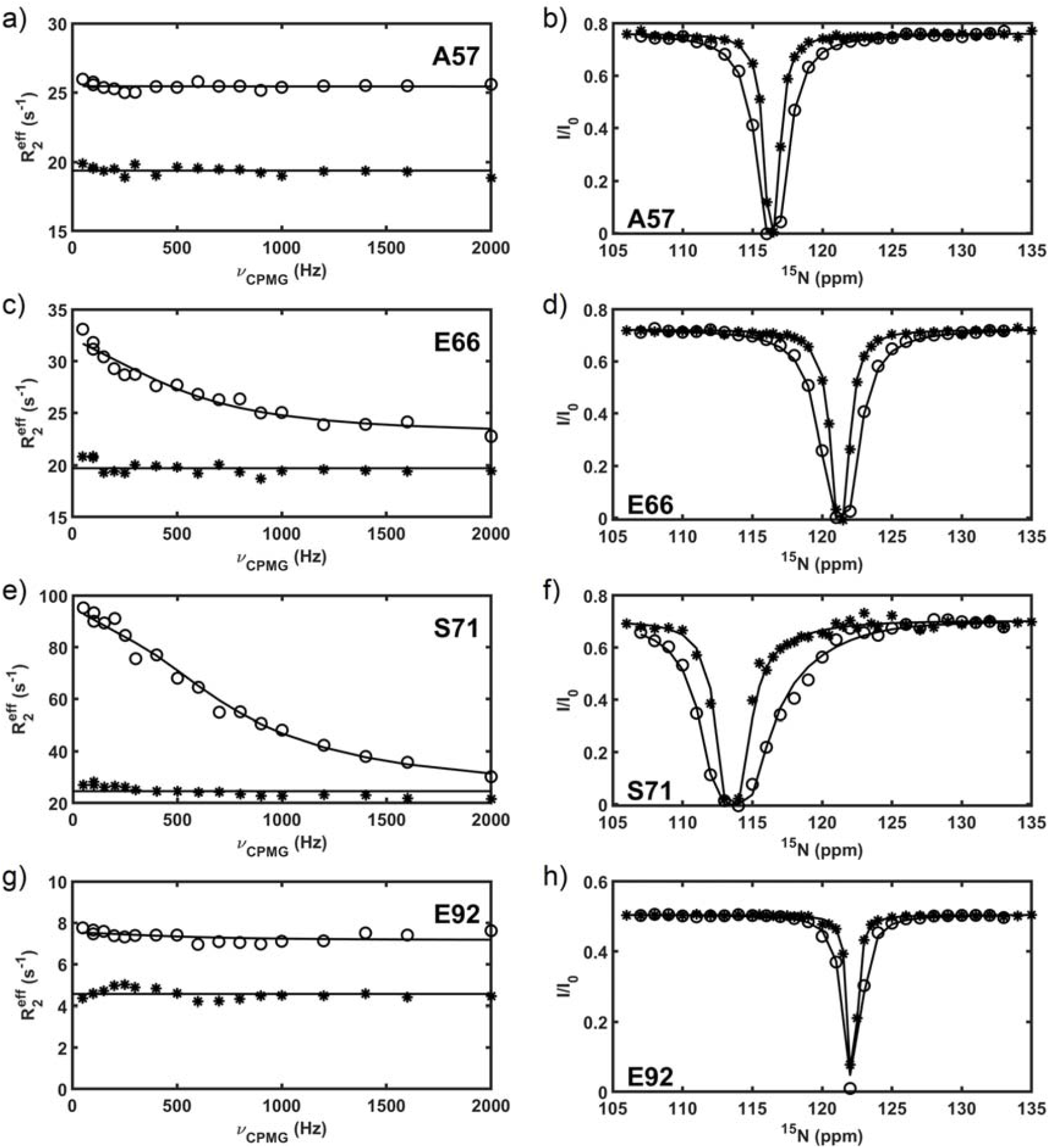
Representative ^15^N relaxation dispersion profiles of DENV2 NS2B residues (a, c, e, and g), with their corresponding CEST profiles (b, d, f, and h). In the CPMG profiles, open circles (o) represent data points collected for the wild-type, while asterisks (*) represent data points collected for the NS2B I73C + NS3pro L115C disulfide bond mutant. In the CEST profiles, open circles (o) represent data points collected on the wild-type sample in the higher saturation field of 26 Hz, while asterisks (*) represent data points collected in the lower saturation field of 13 Hz.

Fitting the data from 13 residues with R_ex_ values larger than 6 s^-1^ globally, we obtained populations of states I and N (pI = 4.0 ± 0.2% and pN = 96.0 ± 0.2%) and the total exchange rate (k_ex_ = 3257 ± 86 s^-1^). ^15^N chemical shifts of state I determined from the data fitting are listed in Table S1. For the other residues with R_ex_ > 3 s^-1^, their chemical shifts in state I were obtained from data analysis by fixing k_ex_, pI and pN at the values derived from the global fitting, which are also listed in Table S1.

### Structure of state I

Past studies have not been able to obtain structural information of the open conformation due to its low population size (∼4% as shown here). In this study, we combined the fitting of both the CPMG and CEST data to obtain both the magnitudes and signs of the chemical shift differences between states N and I, by following the method demonstrated previously(39). From this, ^15^N chemical shifts of the minor state I were calculated and compared with those of both the major state N and unfolded (or disordered) state U (Tab. S1). In terms of ^15^N chemical shifts which are different for folded and disordered proteins, the region of E63–N88 in state I is very similar to that in the disordered state, but significantly different from that in the major native state N (Fig. 4). The result suggests that state I of NS2B adopts a partially unfolded structure: D50–E62 has a native conformation with a regular β-strand (D50–D58), E63–N88 exists in disordered conformations, and E89–R100 remains disordered. In fact, state I resembles the open conformation of NS2B-NS3pro, but it contains no regular secondary structure elements in E63-R100. The conformational exchange between states N and I involves partial unfolding of NS2B and the unfolding rate was estimated to be ∼130 s^-1^ (k_ex_ × pI).

**Figure 4.**
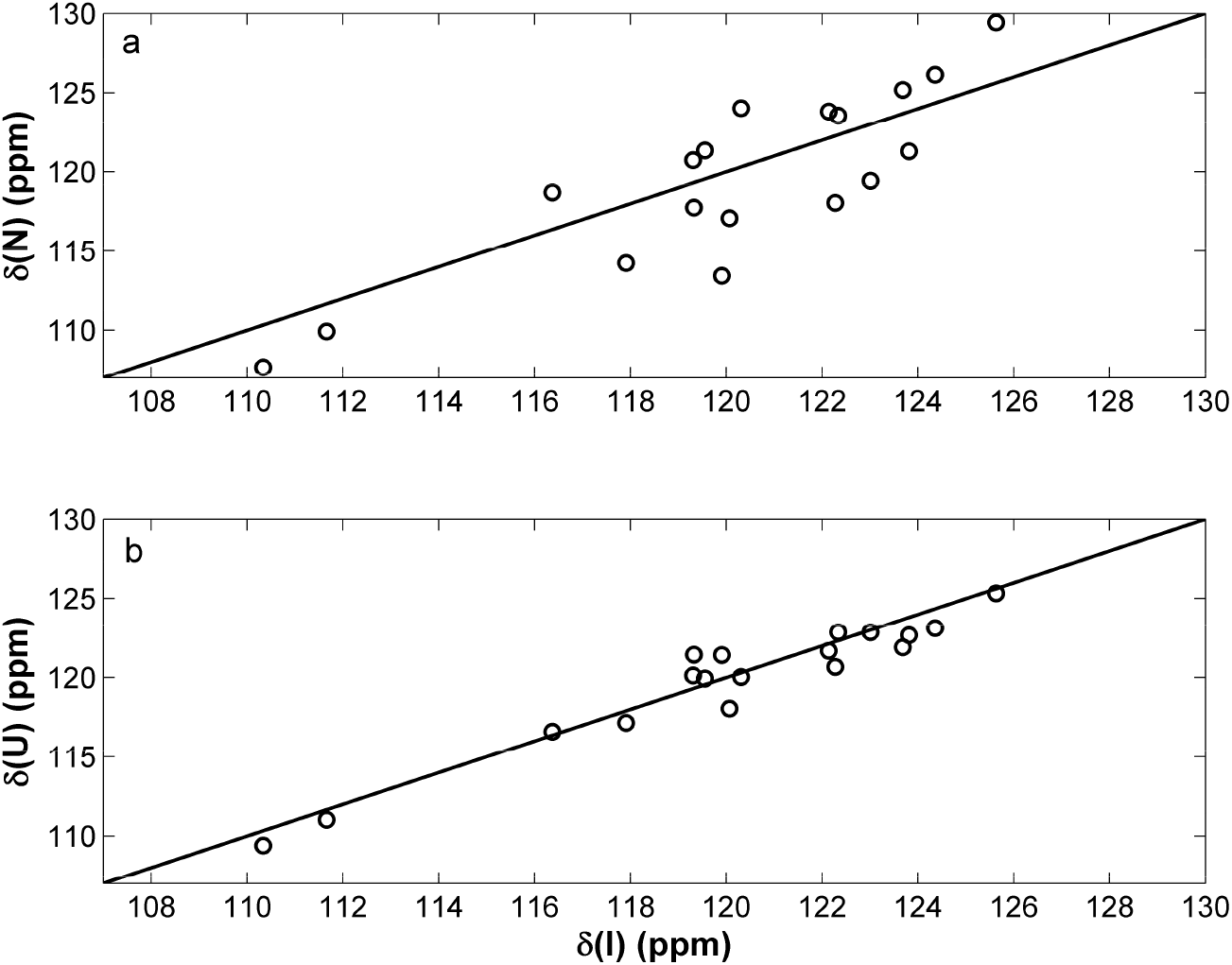
^15^N chemical shift comparisons between the minor state (I) of DENV2 NS2B complexed with NS3pro and its major state (N) (a), and between the state I and the unfolded state (U) (b).

### Effect of mutation on the open conformation population

With the information obtained about the equilibrium between the closed and open conformations of the NS2B-NS3pro complex, we further investigated the effects of perturbing this equilibrium on the activity level of the protease. This was achieved through the generation of point mutants. First, a proline residue was introduced as a single point mutation on the C-terminal β4 of NS2B (M84P) to destabilize the structure and shift the equilibrium towards the open conformation. For the wild type NS2B, the backbone ^1^H-^15^N peaks were well dispersed without significant overlap in the HSQC spectrum (Fig. 1). For the M84P mutant, on the other hand, the correlations became much less dispersed (Fig. 5). This point mutation caused significant changes in peak positions of most residues in the region of E63–N88. Peaks with little chemical shift perturbations belong to the terminal regions (A49–E62, E89–R100) including β1 (Fig. 5). The result suggests that the region of E63–N88 adopts a conformation different from the native conformation. Due to the poor ^1^H chemical shift dispersion characteristic for unfolded protein, this region most likely adopts a disordered form and the mutant mainly exists in an open conformation.

**Figure 5.**
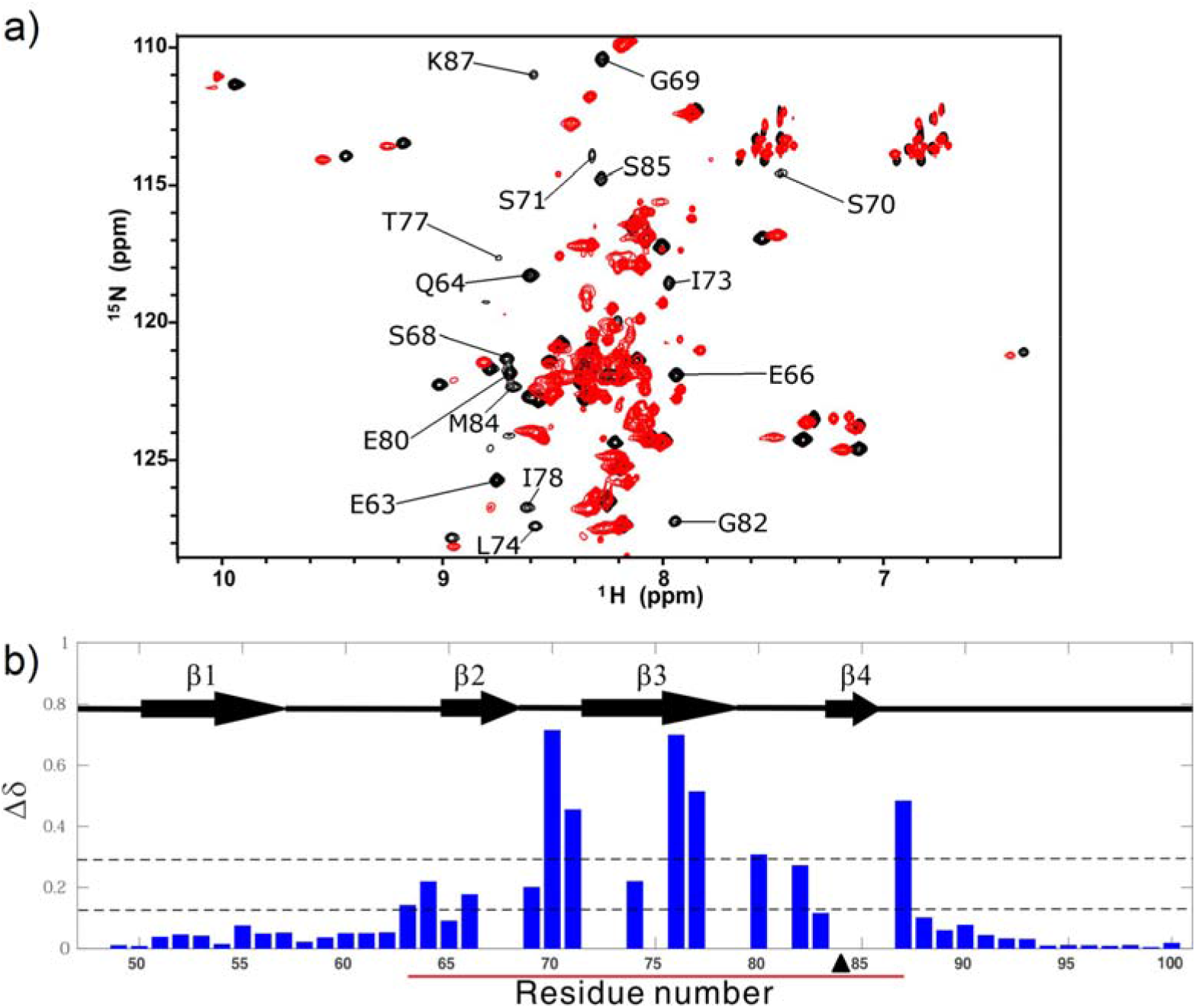
(a) ^1^H-^15^N HSQC spectrum of the M84P mutant (red), overlaid with the ^1^H-^15^N HSQC spectrum of the wild-type DENV2 NS2B-NS3pro (black). Residues with significant chemical shift perturbations are labelled. (b) Combined chemical shift perturbation (Δδ) for each residue of NS2B upon the introduction of the M84P mutation, with the secondary structure of NS2B included within the chart. The region that undergoes conformational exchange in the wild-type is indicated by a red line, while the position of the mutation is indicated by a black triangle. The lower dotted line indicates the average of the chemical shift perturbations, while the higher dotted line indicates the average plus one standard deviation.

To further examine the structural feature of the region of E66–N88 in the M84P mutant, ^15^N HetNOE experiment was performed. For the mutant, the region of G69–M84 was similar to the disordered C-terminal region of N88–L98 with small HetNOE values (<0.25), but the region of D50–Q64 had large HetNOE values (>0.4) (Fig. 2). In comparison with the WT NS2B-NS3pro, the region of G69–M84 in the mutant had significantly smaller HetNOE values and other regions remained nearly unchanged. Because the region of G69–R100 in the mutant is as flexible as the disordered region of 95–R100 in the WT protein, the M84P mutant should be mainly disordered in the region of G69–R100 or exist mainly in an open conformation, consistent with the poor peak dispersion feature. The result indicates that destabilization of β4 by mutation (M84P) shifts the closed-open conformational equilibrium from a closed conformation to an open conformation.

Next, a pair of cysteine residues were introduced as point mutations at both I73 of NS2B and L115 of NS3pro. The cysteine pair is positioned such that the disulfide bond formed under oxidising conditions will lock the C-terminal region of NS2B in place, shifting the equilibrium towards the closed conformation. The HSQC spectra of the WT protein and its disulfide bond mutant (NS2B I73C+NS3pro L115C) were very similar in peak dispersion and positions (Fig. S1a). This mutation caused chemical shift perturbations on a few residues either sequentially or spatially near to the mutation point (Fig. S1b). Additionally, peak intensities were quite uniform, and the peak of residue D81, which was missing in the HSQC spectrum of the wild-type, was visible for the disulfide bond mutant. The disulfide bond mutant had similar HetNOE values to the wild-type (Fig. 2). In fact, loop2 between β2 and β3 became more rigid upon the introduction of the disulfide bond. The results show that the disulfide bond mutant exists mainly in a structure of the state N of the wild-type protein, which is the closed conformation.

To examine the effect of the introduction of the disulfide on the conformational exchange of the complex, RD and CEST experiments were performed. The disulfide bond mutant displayed no relaxation dispersion, even for the residues with large R_ex_ values and far away from I73 in the WT protein (Fig. 3). In addition, the mutant displayed one dip for each ^1^H-^15^N peak. The results show that there are no more conformational exchanges on millisecond and sub-second timescales or the population of the open conformation is too small (<0.5%) to be detected for this mutant. The result also demonstrates that stabilization of the complex shifts the closed-open conformational equilibrium towards the closed conformation by the disulfide bond.

### Effect of mutation on the activity of NS2B-NS3pro

The above results demonstrated that mutations can shift the dynamic equilibrium between the open and closed forms. To see how the change of this equilibrium affects the protease activity, activity assays were performed to compare the mutants with the wild type. The M84P mutant had no activity, while the WT protein was active (Fig. 6 and Tab. 1). The CD spectra of this mutant and the WT protein are very similar (Fig. S2), indicating that the loss of the activity is not caused by the misfolding of the NS3pro but caused by shifting the equilibrium from dominance of the closed NS2B conformation to dominance of the open conformation.

**Table 1.** Kinetic parameters of the mutants compared with the wild-type DENV2 NS2B-NS3pro.

| Construct | $V_{\max}$ (nM/s) | $K_m$ ($\mu\text{M}$) | $k_{\text{cat}}$ ($\text{s}^{-1}$) | $k_{\text{cat}}/K_m$ ($\text{M}^{-1} \text{s}^{-1}$) |
| --- | --- | --- | --- | --- |
| WT | $8.0 \pm 0.2$ | $104 \pm 4$ | $0.80 \pm 0.02$ | $7800 \pm 300$ |
| M84P | - | - | - | - |
| NS2B I73C+<br>NS3pro L115C | $8.3 \pm 0.1$ | $85 \pm 2$ | $0.83 \pm 0.01$ | $9800 \pm 400$ |

-: data not available

**Figure 6.**
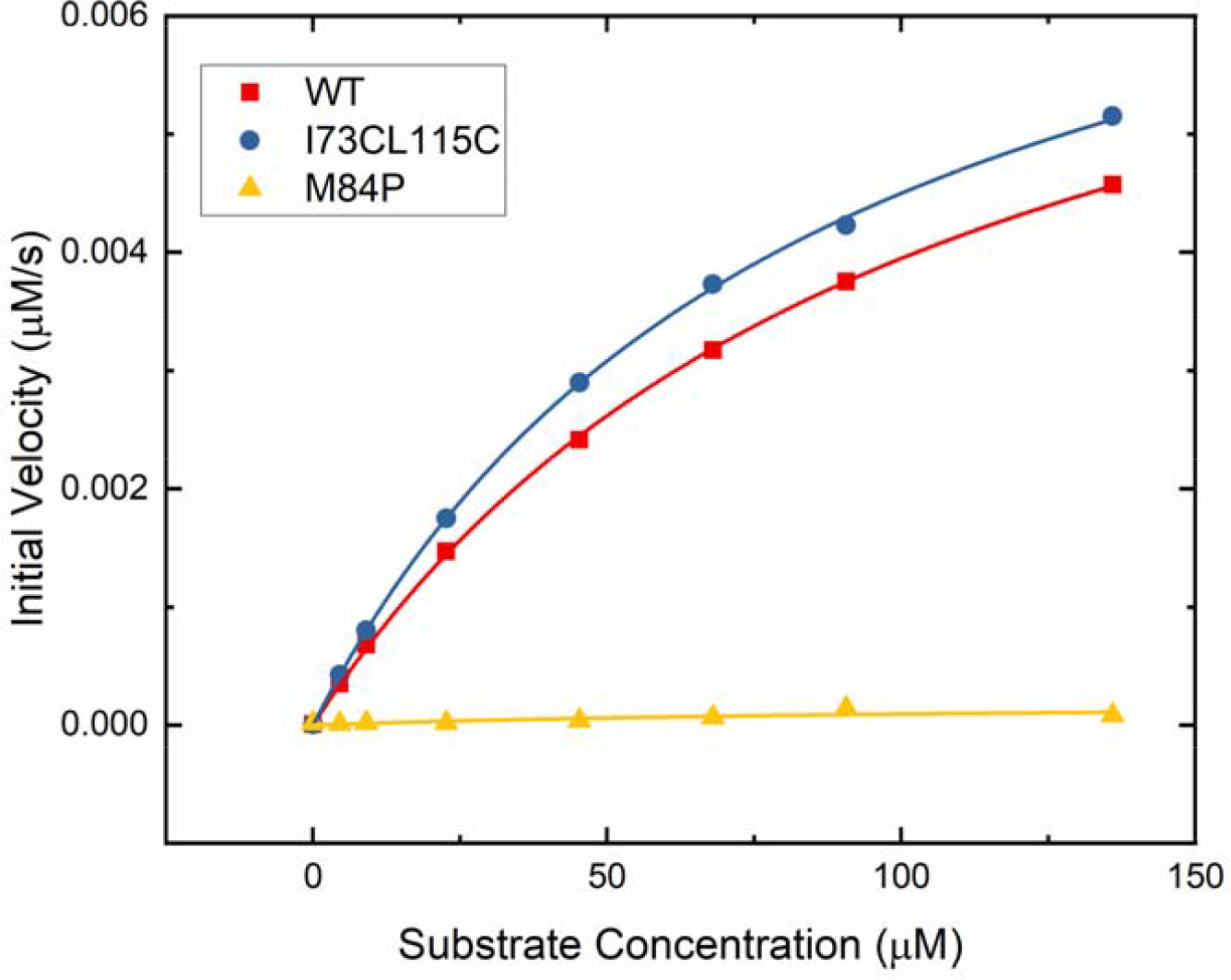
Michaelis-Menten plot for the proline mutant M84P and the disulfide bond mutant NS2B I73C+NS3pro L115C (denoted as I73CL115C), with comparison to the wild-type (wt) DENV2 NS2B-NS3pro.

On the other hand, the disulfide bond mutant NS2B I73C+NS3pro L115C had a slight increase in enzymatic activity (Fig. 6 and Tab. 1). As the population size of the closed conformation increased by only 4% by introduction of the disulfide bond, the enzymatic activity enhancement should be insignificant by stabilization of the wild type NS2B-NS3pro complex.

## Discussion

The existence of the open conformation of the NS2B-NS3pro complex has baffled researchers since its discovery in 2006 (18). It was initially thought that the dengue NS2B-NS3pro complex is predominantly in the open conformation, in contrast to the West Nile NS2B-NS3pro. This was due to the lack of a crystal structure showing dengue NS2B-NS3pro in the closed conformation. This was further supported by some NMR studies, which suggested that the C-terminal region of NS2B is dissociated from NS3pro most of the time(40, 41). This has been changed recently when a crystal structure of dengue NS2B-NS3pro was solved in the presence of a peptide inhibitor(20). Recent NMR studies have also given an estimate of not more than 5% of the complex in the open conformation(38, 42). Various crystal structures(18, 20, 21, 23-26) and NMR studies (38, 42, 43) have provided evidence to support that the closed conformation is the dominant and enzymatically active form, and therefore biologically relevant. Our results are in agreement with these studies, showing that the dengue NS2B-NS3pro complex is mainly in the closed conformation (pN = 96%), which is the active form.

Having the open conformation as an alternative inactive form seems pointless, but for the open conformation to persist through the selective pressures of evolution, it must play a significant role in nature. To understand its role in the protease function, structural information of the open conformation is required. Previous attempts to study the open conformation have been unsuccessful due to its small population size, as well as severe NMR peak broadening and overlapping(28, 38, 42). In this study, we overcame these problems using two approaches. First, we expressed NS2B and NS3pro sequentially such that a complex between isotopically labelled NS2B and unlabelled NS3pro was obtained. This has reduced the number of NMR peaks on the HSQC spectrum from at least 200 to around 50, reducing the amount of peak overlapping. This allowed us to achieve ∼95% amide peak assignments for NS2B (Fig. 1). Next, we performed a combined fitting of both the CPMG and CEST data to a two-state model to obtain the chemical shifts of the open conformation (Tab. S1). These allowed us to obtain structural information of the open conformation, which revealed that the C-terminal region of NS2B is unfolded or disordered in the open conformation of NS2B-NS3pro (Fig. 4). The open conformation obtained here is in contrast to some of the crystal structures which suggests the existence of the β-hairpin in the open conformation(21, 22). The β-hairpin structure is likely to be an artefact that occurred during the crystallisation process. Better understanding of the functional role of the open conformation needs further studies on structures, dynamics and protease activities of NS2B-NS3pro complexes from other dengue virus serotypes.

Our study has revealed through activity assays of various mutants that shifting the closed-open conformational equilibrium to either side causes an effect on the proteolytic activity of the protease (Fig. 6, Tab. 1). This shifting can occur either through natural means like pH and salt concentration(42), or artificial means like site-directed mutagenesis and small molecule inhibitors (44, 45). The protease activity can therefore be controlled through this equilibrium by both nature and man. From the perspective of nature, it could be used by the virus as an auto-regulatory mechanism to tune down its protease activity when the environmental conditions are not favourable for its replication. On the other hand, from the perspective of man, the equilibrium could be exploited during drug design to inhibit the viral protease. Although the closed conformation is more biologically relevant, the open conformation may be the key to developing drugs with great efficacy and specificity to inhibit the dengue protease.

## Conclusion

Our results show that the dengue NS2B-NS3pro complex is predominantly in the closed conformation and the open conformation is unfolded in region E63–K87 of NS2B. The shifting of the equilibrium between the open and closed conformations has also been shown to change the activity level of the protease. This closed-open conformational equilibrium is possibly used by the virus for self-regulation, and at the same time can be exploited in future drug design to treat dengue.

## Acknowledgments

This research was supported by grants from Singapore Ministry of Education (Academic Research Fund Tier 3, MOE2012-T3-1-008; Academic Research Fund Tier 1, R-154-000-C03-114).

## Author Contributions

DY conceived the project. WHL, WL, and JSF performed experiments and data analysis. WHL and DY wrote the manuscript with input from all authors.

